# *De Novo* Prediction of RNA 3D Structures with Deep Learning

**DOI:** 10.1101/2021.08.30.458226

**Authors:** Julius Ramakers, Christopher Frederik Blum, Sabrina König, Stefan Harmeling, Markus Kollmann

## Abstract

We present a Deep Learning approach to predict 3D folding structures of RNAs from their nucleic acid sequence. Our approach combines an autoregressive Deep Generative Model, Monte Carlo Tree Search, and a Score Model to find and rank the most likely folding structures for a given RNA sequence. We confirm the predictive power of our approach by setting new benchmarks for some longer sequences in a simulated blind test of the RNA Puzzles prediction challenge.

## Main

The three-dimensional (3D) folding structure of RNA can have significant impact on its role as a mediator and modulator of genetic information. Such structure related effects become most apparent for synthetic RNA, where changes in the secondary and tertiary conformations can significantly alter RNA stability, translational/editing efficiencies, and binding affinities in case of aptamers ^[2]^. Consequently, the automated prediction and the targeted design of RNA tertiary structures would be an important step to improve the functionality of RNA, in particular for RNA therapeutics ^[1,3]^.

Algorithms for predicting RNA 3D structure from nucleotide sequence ^[4]^ are dominated by three approaches: (i) template based methods ^[5]^, which decompose known structures into 1-to 3-mer fragments and combinatorially reassemble them to find the structures with lowest molecular interaction energies ^[6]^, (ii) coarse grained force field methods that minimise interaction energy by stochastically displacing groups of atoms ^[7]^, and (iii) comparative modelling methods that are based on the availability of homologous structures. Despite the steady increase in affordable computing power and the use of more accurate energy functions [6], the *de novo* structure prediction of larger RNAs (> 80 nt) still remains challenging ^[5]^.

For proteins, the benchmark for predicting 3D structures with atomic resolution is set by deep learning approaches that take sequence information as input and predict both the distances between *C_α_* or *C_β_* atoms and the dihedral angles to determine the conformation of the polypeptide backbone ^[8,9]^. The crucial input information comes from homologs found by multiple sequence alignments (MSA) that carry information about the global folding structure – in particular information for identifying residues that are in contact ^[10]^. These global structural constraints can be inferred from correlations between amino acid substitution frequencies that arise from an evolutionary selection pressure for stably folded protein structures ^[11]^. The ability of deep neural networks to learn complex statistical patterns in high dimensional spaces and to generalise well across training examples makes deep learning approaches conceptually attractive for predicting protein structures, if the number of homologous sequences and the number of independent training examples is sufficiently large ^[12]^.

Transfering deep learning approaches from protein to RNA is difficult as they differ significantly in the molecular structure of their residues, which strongly affects their folding principles. First, in contrast to almost all proteins, RNAs can fold into different alternative structures under physiological conditions that are either stable or visited over time with high probability ^[13]^. Second, accurate prediction of the conformation of the RNA main chain (phosphate backbone) is insufficient to predict the position of the side chains (nucleobases). This difference to proteins arises from the fact that the secondary structure of proteins is determined by hydrogen bonds within the peptide backbone, whereas the secondary structure of RNA is determined by hydrogen bonds between the side chains. Third, training of deep learning models requires a large amount of independent training examples, but the protein data bank (PDB) contains two orders of magnitude less experimentally determined RNA structures than protein structures. Finally, the less conserved RNA structures make it much harder to identify homologs for MSA and therefore the crucial information about global folding constraints is in many cases not accessible. On the contrary, there exist structural probing methods to estimate the probability of each nucleotide to be part a of a base paring interaction, such as SHAPE^[14]^ or DMS^[15]^. However, unlike MSAs, structural probing methods can only give an estimate if a nucleotide is in contact, but lack direct information about the contact parter. Moreover, structural probing data represents an ensemble average over the structural conformations that a given RNA can take and therefore can provide only useful information if the secondary structure is sufficiently stable.

In our deep learning approach included some best practices for training deep neural networks. First, deep neural networks strongly benefit from end-to-end learning, where gradients for updating parameters are allowed to propagate from the objective function back to the input, thereby avoiding extensive preprocessing steps that might reduce the information content ^[16]^. Second, the inductive bias induced by the network architecture should match the structure of the data. We therefore combined self-attention layers (Supplementary Information) to extract long-range correlations within the RNA sequence and used convolutional layers to predict local correlations in the RNA 3D structure ^[17,18]^. Third, the final performance of a deep learning model depends significantly on (i) the neural network size, (ii) the amount of training data, and (iii) the training time. The generic empirical observation, which is also confirmed in this work, is that increasing (i)-(iii) increases the prediction accuracy ^[19]^. Consequently, we used advanced data augmentation techniques, which allowed us to train larger networks that were able to model more complex mappings and achieve better generalisation (Methods). To augment the 788 RNA only structures extracted from the PDB, we generated substructures of length *L* = 100 nt by randomly cropping the 108 PDB structures with length *L* > 100 nt. As the cropped substructures are in contact with the remaining part of the structure, we used binary indicator variables to mark all nucleotide pairs that are in contact with the remaining structure and showed the corresponding distance classes (Fig. 1) at the input during training. As these substructures made up most of the training set, the model effectively learned to predict substructures that were constrained by the remaining part of the RNA structure. To predict free folding RNA structures we employed a transfer learning approach by taking the prediction model for the substructures and set the binary indicator variables to zero. We fixed the length of the substructures to 100 nt, as shorter substructures have more contacts on average, which makes transfer learning for free folding structures less reliable.

**Figure 1.**
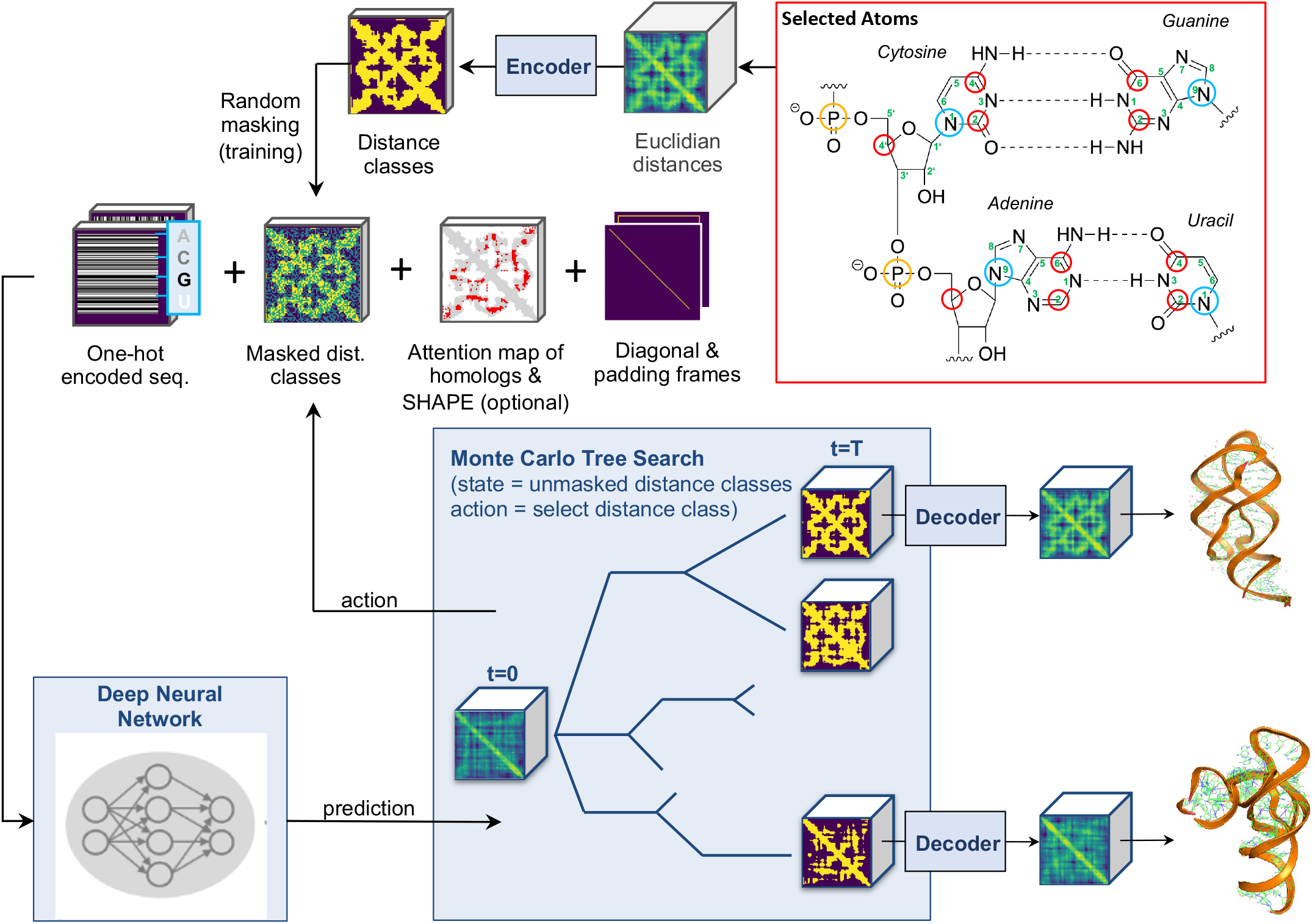
Data flowchart for the RNA structure generation process. The PDB structure is represented by Euclidian distances between nucleotide pairs, where the position of each nucleotide is determined by five selected atoms. The resulting 5 × 5 Euclidian distances for each each nucleotide pair are encoded into *K* discrete distance classes by a VQ-VAE ^[20]^. The generation process uses a Deep Neural Network (DNN) to predict probability values for the distances classes. From these predictions a single distance class for a single nucleotide pair is selected according to the MCTS policy (Methods) to iteratively generate a path in the search tree. At each iteration the currently selected distance classes and the sequence information are presented as input to the DNN. Once all distance classes are selected, the Euclidian distances can be recovered by the VQ-VAE decoder. A Score Model (Methods) selects the most promising generated structures, which are then further refined by minimising a coarse grained molecular energy function^[7]^.

To encode 3D RNA structures, we used a rotational invariant representation that was given by the Euclidian distances between nucleotides, with each nucleotide position uniquely determined by a set of 5 selected atoms, where different sets were taken for purines and pyrimidines (Fig. 1). We made use of a Vector Quantised Variational Autoencoder^[20]^ (VQ-VAE) to compress the 5 × 5 Euclidian distances between the selected atoms into *K* classes for each possible nucleotide pair. We refer to these classes as distance classes, as the *K* = 3 classes we used throughout this work agree well with the qualitative distance measures “near”, “intermediate”, and “far” (Supplementary Information). The encoded distance classes represent the targets used for training a Deep Generative Model that takes sequence information and masked targets as input. The task of the Generative Model was to predict the probabilities of the masked distance classes. For the masking, we first selected the fraction of nucleotide pairs (pixels) to be masked by randomly drawing an integer number *n* from the set {1, 2, .., *L*^2^}, with *L* the sequence length, and then randomly selecting *n* out of *L*^2^ pixels whose one-hot encoded target values were then overwritten by assigning each distance class the same value. Training neural network architectures on such “gap-filling” tasks shows surprisingly strong generalisation behaviour and has resulted in state-of-the-art results for learning words representation in Natural Language Processing (NLP) and for image generation in computer vision ^[21,22]^. After training, a structure can be iteratively built up by sampling a distance class for each nucleotide pair according to a MCTS search algorithm (Methods) and presenting the selected distance class at the input (Fig. 1). Although our generative model allows to estimate the likelihood for each predicted structure by making use of the chain rule for probability mass functions ^[22]^, this value is in general unreliable ^[23]^. We therefore trained a Score Model (Methods) that allowed to score the match between sequence and generated structures, similar to a value function in reinforcement learning ^[24]^. Each predicted, one-hot encoded distance matrix with high score was mapped back to an Euclidian distance matrix, using the decoder of the VQ-VAE. The Euclidian distances were further fine tuned by minimising a coarse grained, physical RNA energy function ^[7]^.

We first tested the reconstruction accuracy of the VQ-VAE as a function of the number of distance classes (Fig. 2a). For *K* = 8 classes, the reconstruction error approached the average experimental resolution of 2.8 Å root-mean-square error (RMSE). For *K* = 3 classes the median reconstruction error was still in the range of the best predictions with 4 Å RMSE. Next, we simulated a blind test by evaluating the accuracy on a held out test set that is given by the crystal structures of the RNA puzzles challenges ^[25]^ (Fig. 2b,c), including only free folding RNA structures. We observed significant improvements for RNA structure prediction problems that were classified as difficult ^[5]^ (RNA-Puzzles 5 and 7) due to their longer sequence (> 100 nt) and their lack of homology to known structures (Fig. 2d).

**Figure 2.**
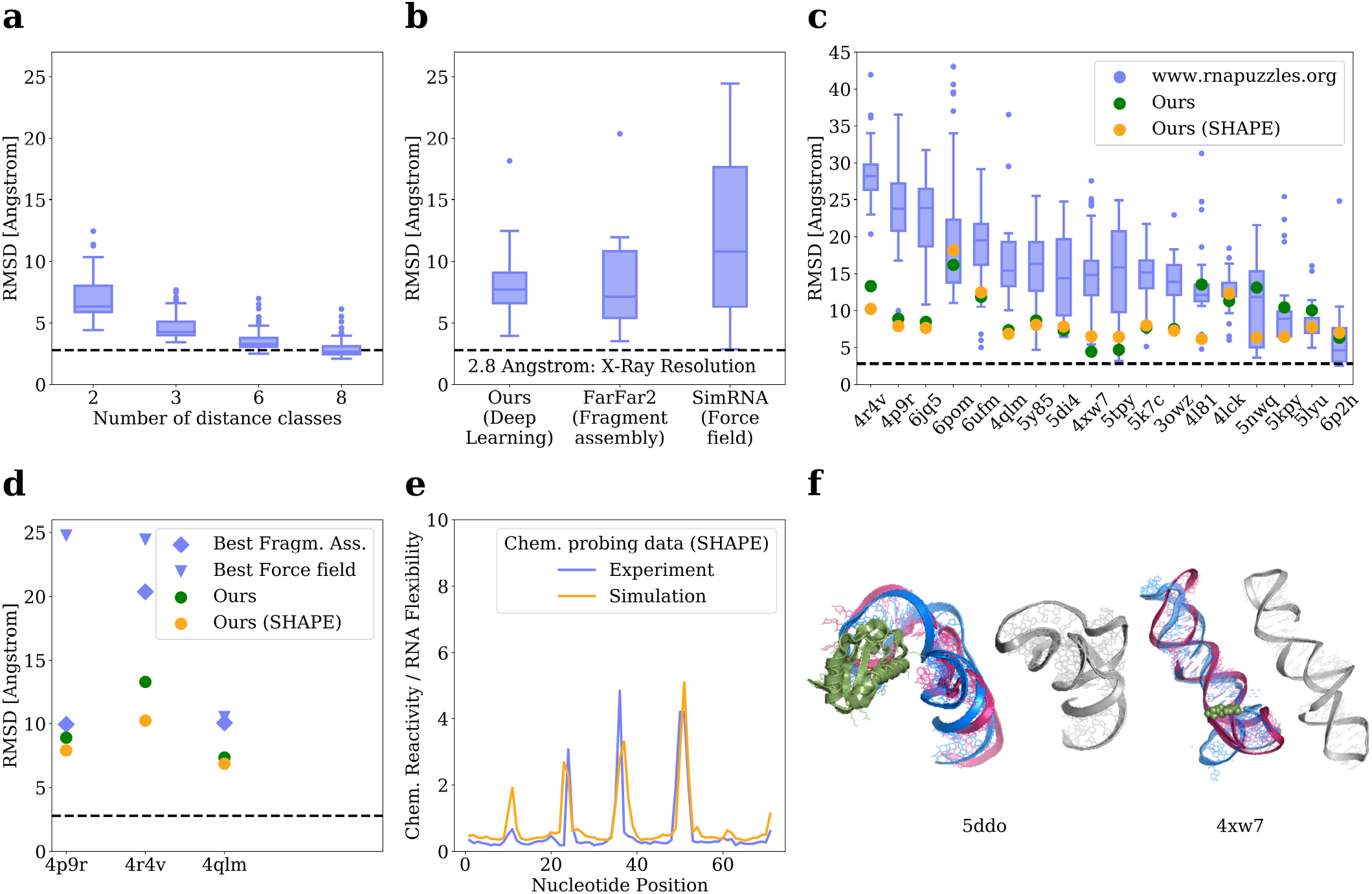
Evaluation of the structural predictions. **a,** Reconstruction error resulting from encoding and decoding RNA 3D structures of the test set as a function of the number of distance classes, **b,c** Simulated blind tests of the RNA Puzzle Challenges in comparison to other approaches, with or without simulated SHAPE reactivity data and homologous sequences as additional input. The predictions are sorted in descending order of the sequence length (left to right) **e,** Reconstruction error of longer RNA puzzles that lack both structural and sequence homology (PDB 4P9R: 189 nt, 1 homolog; PDB 4R4V: 185 nt, 2 homologs) in comparison to (PDB 4QLM: 108 nt, 172 homologs) that lacks only structural homology. RMSE over the first 100nt **e,** Simulated chemical probing data in comparison with experimentally measured reactivities^[26]^ (SHAPE) for PDB 1Y26, **f,** Most likely alternative structures, as predicted by the Score Model, for the Gluatamine riboswitch and the ZMP riboswitch (Supplementary Fig. 3)

To investigate the effect of structural probing data (SHAPE), we used a force field model ^[7]^ to simulate SHAPE reactivities which are shown at the input during training. We thereby assumed that single stranded RNA is more flexible than double stranded RNA and thus shows higher mean squared displacement (MSD) of atoms during the force field simulations. The simulated MSD values show good agreement with the experimentally determined SHAPE reactivities (Fig. 2e). We observed a small improvement in prediction accuracy on average when we presented both SHAPE data and MSAs of homologous sequences at the input, which indicates that the additional constraints imposed by simulated SHAPE data and evolutionarily constrained nucleotide-nucleotide interactions provide only little additional information to our model for RNAs of length *L* ≤ 100 nt. However, we found that this additional input data allowed to infer global structural information (Supplementary Fig. 2), thereby accelerating MCTS. This acceleration might become crucial in cases where exploring the global structural space by MCTS is the limiting factor.

The ability of our approach to find alternative structures can be used to predict the different states of riboswitches (Fig. 2f). We simulated a blind prediction test by removing all homologous structures from the training set for the Glutamine Riboswitch (PDB: 5*DDO*) and the ZMP Ribosowitch (PDB: 4*XW* 7). The predicted alternative structures, which represent two highest ranked branches of the MCTS by the Score Model, confirm the general viewpoint that riboswitches work by a ligand mediated stabilisation of one structural conformation.

## Acknowledgements

We acknowledge funding from the Jürgen Manchot Foundation and the CEPLAS - Cluster of Excellence on Plant Sciences (EXC 2048). We further acknowledge the computing time granted by High-Performance Computing Centre at the Heinrich Heine University Düsseldorf. The calculations for this research were conducted with computing resources under the project DeepGenome. We especially thank the HPC team for their technical support.

## Online Methods

### Data extraction and preprocessing

We extracted 1183 RNA single chain structures from the Protein Data Bank (PDB) that correspond to 788 unique RNA sequences of length *L* ≥ 20 nt, with some structures measured under different experimental protocols. RNA structures in complex with protein/DNA were discarded. The extracted structures were grouped according to their sequence similarity, using hierarchical clustering with a similarity cutoff of 0.7. We split the clusters into training, development, and test sets (653, 76, and 59 sequences, respectively). To augment the structural data, we carried out Molecular Dynamics (MD) simulations^[7]^ for each of the 788 sequences that were initialised by the atom positions of structural variants that correspond to the same PDB id (NMR ensembles or symmetrical copies of biological assemblies). We ran the simulations independently 50 times with a varying number of time steps, so that we obtained “drifted structures” at 1, 3 and 5 Å root-mean-square error (RMSE) from the original PDB structure. We generated additional sequences by randomly cropping the unique RNA sequences of length *L* ≥ 100 to length *L* = 100 nt. For each of the 8048 resulting sequences with length *L* ≤ 100, we selected the corresponding substructures from the MD simulations. We memorised the nucleotides that were in contact with the remaining structure using a distance cutoff of 3.3 Å . After data augmentation, the training, development, and test sets contained 352350, 28050, and 22000 structures, respectively. We determined the position of each nucleotide by 5 selected atoms (Fig. 1) and compressed the 25 possible real distances between the selected atoms for any nucleotide pair into *K* = 3 distance classes using a Vector Quantised Variational Autoencoder (VQ-VAE).

### Autoregressive Generative Model

The generation of a 3D structure, ***s***, from sequence information, ***x***, was carried out iteratively by first selecting a nucleotide pair (pixel) with index *i* ∈ {1, .., *N*} from the *N* = *L*(*L* − 1)/2 possible pairings and subsequently selecting a distance class *k_i_* ∈ {1, .., *K*} according to the class probabilities predicted by the Generative Model, *P* (***k***|***s****_t_,* ***x***). The selected distance class was then one-hot encoded, resulting in an updated input structure ***s***_*t*+1_ ← ***s****_t_*. We denote by 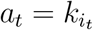 the “action” for the *t*-th iterative step in the generation process and defined the current structural state by ***s****_t_* = (*a_t_*, …, *a*_1_). Actions are never overwritten during the generation process, which starts from an empty set of actions ***s***_0_ by “masking” all pixels as defined below. The Generative Model was realised by a feed-forward neural network *P* (***k***|***s****_t_,* ***x***) = Π*_i_ P_i_*(*k_i_*|***s****_t_,* ***x***) that predicted probabilities for all distance classes ***k*** = (*k*_1_, *k*_2_, …, *k_N_*) in parallel, given the previous actions ***s****_t_*. The conditional independence of the predicted class probabilities is a consequence of the deterministic network architecture, where the outputs (class probabilities for each pixel) are uniquely determined by the input. The complete input feature map of the autoregressive model comprised of (i) a one-hot encoding of the 16 possible nucleotide pairs (AA,AC, …, UU) for each pixel, (ii) the already set distance classes in the autoregressive process, ***s****_t_*, (iii) coordinate frames that included the diagonal as symmetry axis and padded regions for sequences of length *L* < 100, (iv) the output of a Self-Attention layer ^[18]^ that takes structural probing data and homologous sequences as input (Fig. 1). Training was carried out by showing for ***s****_t_* a random fraction of distance classes (masked target) at the input, where the one-hot encoding for the *K* distances classes was substituted by information that was related to the target logits, e.g. (1, −1, −1) if the target was the first distance class and (0, 0, 0) if the target class was masked. The number of masked distance classes shown at the input was distributed according to a truncated half-normal distribution, to enforce that almost complete targets are shown less frequently during training. The feed-forward network architecture was a residual network with 16 residual blocks and 26 channels in each hidden layer, trained by early stopping (Supplementary Information). The generative model allowed to compute likelihood estimates of a structure ***s****_N_* for a given nucleotide sequence ***x*** by making use of the chain rule of probability, 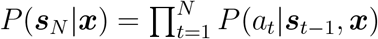.

### Score Model

The likelihood estimate was improved by learning a Score Model *D*(***s****_N_*, ***s***′_*N*_; ***x***) (discriminator) that was trained to distinguish between correct and incorrect structures-sequence pairs by maximising the objective

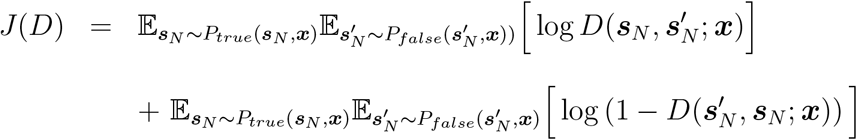

with *D* defined by

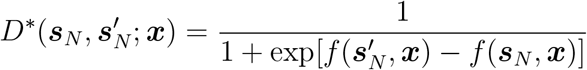

and *f* (***s***′_*N*_, ***x***) being the scalar output of a deep neural network. The theoretically optimal solution 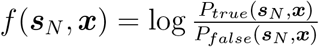 is typically not reached by optimisers based on stochastic gradient decent ^[27]^. Here, “true” corresponds to original PDB examples and “false” to PDB examples with drifted atom position using Molecular Dynamics Simulations ^[7]^ under high temperature and encoded by the VQ-VAE or structures predicted from the Generative Model. The discriminator compares two complete structures with respect to their match to a given sequence, which is in contrast to the absolute likelihood estimate from the chain rule of probability. The value *f* (***s***′_*N*_, ***x***) is used to rank the predictions that are sampled from the Generative Model. For the Score Model, we used a Residual Network architecture with 8 residual blocks, where blocks were connected by down-sampling layers using stride 2 convolutions (Supplementary Information).

### Structural Sampling

Unlike proteins, RNAs frequently fold into different structures under physiological conditions. To identify the structures that occur with high probability, we had to sample the large combinatorial space of possible structures and rank them according to their corresponding likelihood. For RNAs of length *L* = 100 nucleotides, the combinatorial space of allowed distances is given by *K*^*L*(*L*−1)/2^ > 10^2000^ for *K* ≥ 3 and thus exceeds the number of possible games, ~ 10^700^, that can played in the board game Go. To realise fast sampling, we borrowed search strategies from reinforcement learning (RL). We aimed to find the best sequential ordering (*a_t_*, …, *a*_1_) for presenting distance classes at the input, such that after a minimum number of autoregressive steps the probability masses for the remaining nucleotide distances became highly concentrated into one class. This ordering allowed to predict the final structure after *T* ≪ *N* steps by selecting the most likely distance classes for the set of remaining masked pixels, *M_T_*, in parallel

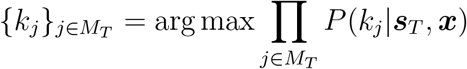

To find a close to optimal autoregressive ordering, we developed a variant of the Monte Carlo Tree Search (MCTS) that accounts for the fact that some actions affect the global structure more that others. For a given nucleotide sequence, a tree of possible RNA structures can be built iteratively by connecting incomplete structures (nodes), *s_t_*, with actions, *a_t_*, such that each leaf node *s_L_* of the current tree can be reached by a unique path of actions *s_L_* = (*a_L_, …, a*_1_). For selecting the actions to reach a leaf node from the root node (empty set of actions), *s*_0_, we followed the selection rule (policy)^[28]^

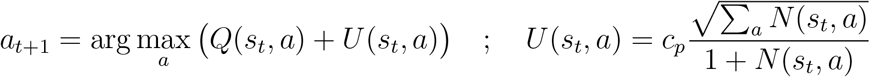

with *Q*(*s_t_, a*) the expected entropy reduction of *P* (*a*|***s****_t_,* ***x***) if action *a* is taken, *N* (*s_t_*, *a*) a counter how often actions that connect the root node with a leaf node pass through the state-action pair (*s_t_, a*) under the actual policy, and *U* (*s_t_*, *a*) a term that upweights rarely visited actions (exploration), with *c_p_* being a tuneable constant. After reaching a leaf node, *s_L_*, the tree is expanded by randomly selecting a subset *S_R_* of the remaining masked pixels, and from *S_R_* a subset *S_H_* ⊂ *S_R_* of actions that result in sufficiently strong entropy reduction Δ*H_L_ > λ* log *K* for the predicted class probabilities, with *λ* = 1 and

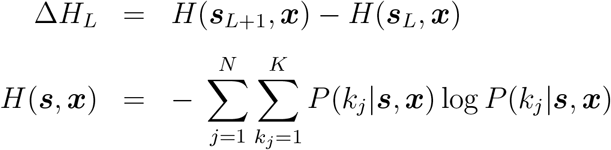

For |*S_H_* | = ∅ the leaf node *s_L_* is added to set of terminal nodes *S_T_* ← *S_T_* ∪ *s_L_* and for |*S_H_* | = ∅ the tree is enlarged by |*S_H_* | nodes in parallel, initialising *N* (*s_L_*, *a*) = 1 and *Q*(*s_L_*, *a*) = *υ*(***s***_*L*+1_|***x***) for all *a* ∈ *S_H_*, with value function the entropy reduction rate

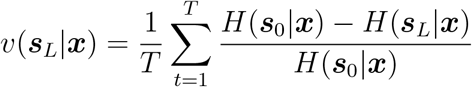

Along the path of actions from the root node to *s_L_* = (*a*_1_, .., *a_L_*), we updated for all *t* ∈ {1, .., *L*} the visit count *N* (*s_t_*, *a_t_*) ← *N* (*s_t_*, *a_t_*) + |*S*| and subsequently the expected reward 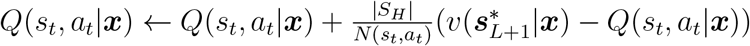, using the ‘winner takes it all’ value function *υ*(***s***\*_*L*_|***x***), with 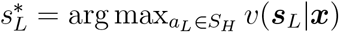. For leaf nodes that were terminal nodes, *s_L_* ∈ *S_T_*, we updated *N* (*s_t_, a_t_*) ← *N* (*s_t_, a_t_*) + *N_expl_*, with *N_expl_* = 10 a hyperparameter, along the path and leave *Q*(*s_t_, a_t_*) unchanged to enforce exploration of different structures.

### Structural Ensemble

We generated complete structures for the set of terminal leaf nodes *S_T_* by arg max sampling from *P* (***k***|***s****_T_*, ***x***), as introduced above. We ranked the complete structures relative to each other according to the discriminator output. The resulting ensemble was a subset of the possible structures that an RNA can take. As we started out to find the find the most likely structure by MCTS, with alternative structures as a by-product of the search, the resulting ensemble was highly biased towards structures with high likelihood.

## Supplementary Information

- Data Preparation and Preprocessing
- Deep Learning Architectures and Training Hyperparameters
- Supplementary Figure 1: Generator Network: Encoding of Sequence Information.
- Supplementary Figure 2: Attention: Improved performance from adding Structural Probing data (SHAPE) and homologous sequences.
- Supplementary Figure 3: MCTS: Search tree with two distinct leafs.

### Data Preparation and Preprocessing

#### Source data

We used only the publicly available RNA 3D structures from the protein data bank (PDB) for both training and validation. We carried out several steps to select adequate structures based on the information in the PDB entry type file^1^ (pdb entry type.txt) and the PDB files themselves. In particular, we only kept a structure if (i) it had only RNA chains (entry-type in pdb entry type.txt was “nuc”), (ii) the resolution method was either X-ray diffraction or NMR (”method” in pdb entry type.txt was either “diffration” or “NMR”), (iii) the PDB structure was downloadable^2^, (iv) the “resNames” were either A, C, G or U, (v) the chain had at least 14 nucleotides, or at least 7 nucleotides if the structure was composed of multiple chains, (vi) for structures with 3 or more chains, all chains were given by unique sequences.

#### Neural Network input data format

All PDB structures were converted into a reduced 5-atom positional representation for each residue^[7]^ (Fig. 1). For Guanosine monophosphate and Adenosine monophosphates, we used the P, C4’, C2, C6 and N9 atoms. For Cytosine monophosphates and Uridine monophosphates, we used the P, C4’, C2, C4 and N1 atoms. To distinguish between purine and pyrimidine residues (G/A and C/U, respectively), we encoded the cartesian coordinates (*x*, *y* and *z*) of the 5 atoms by an 8 × 3 coordinate matrix and indicated the valid atoms (rows) by an 8 × 1 mask array (Supplementary Table 1).

**Supplementary Table 1.**
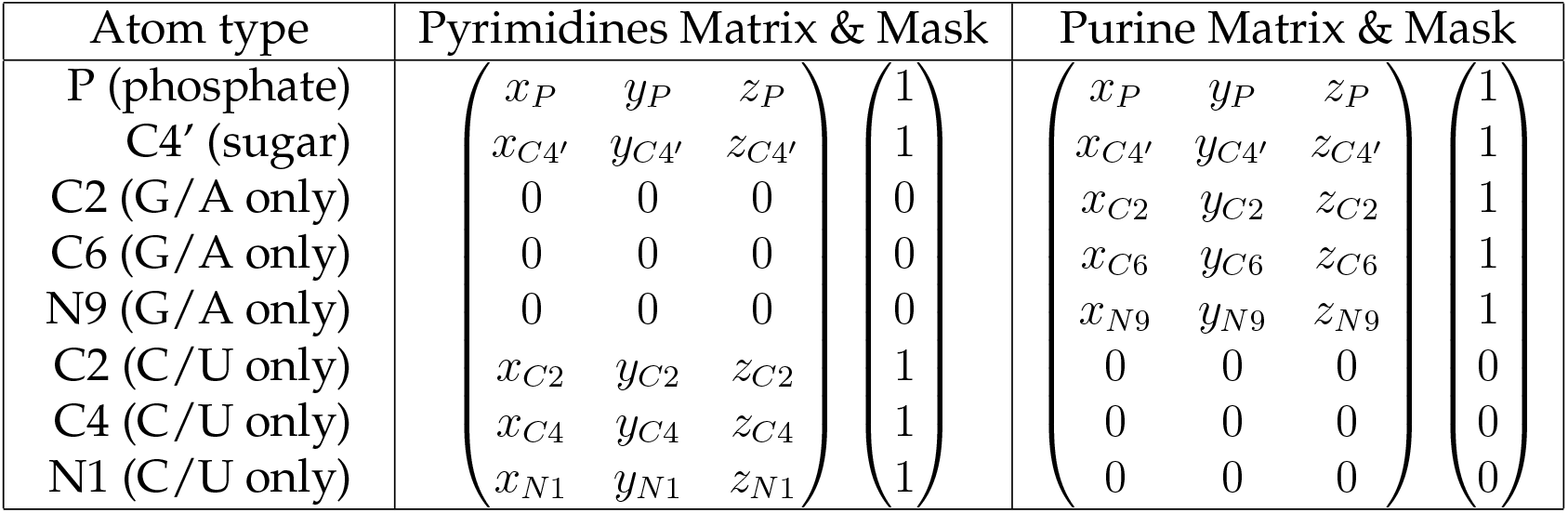
Encoding of purine (G/A) and pyrimidine (C/U) coordinates.

| Atom type | Pyrimidines Matrix & Mask | Purine Matrix & Mask |
| --- | --- | --- |
| P (phosphate) | $\begin{pmatrix} x_P & y_P & z_P \\ x_{C4'} & y_{C4'} & z_{C4'} \\ 0 & 0 & 0 \\ 0 & 0 & 0 \\ 0 & 0 & 0 \end{pmatrix} \begin{pmatrix} 1 \\ 1 \\ 0 \\ 0 \\ 0 \end{pmatrix}$ | $\begin{pmatrix} x_P & y_P & z_P \\ x_{C4'} & y_{C4'} & z_{C4'} \\ x_{C2} & y_{C2} & z_{C2} \\ x_{C6} & y_{C6} & z_{C6} \\ x_{N9} & y_{N9} & z_{N9} \end{pmatrix} \begin{pmatrix} 1 \\ 1 \\ 1 \\ 1 \\ 1 \end{pmatrix}$ |
| C4' (sugar) |  |  |
| C2 (G/A only) |  |  |
| C6 (G/A only) |  |  |
| N9 (G/A only) |  |  |
| C2 (C/U only) | $\begin{pmatrix} x_{C2} & y_{C2} & z_{C2} \\ x_{C4} & y_{C4} & z_{C4} \\ x_{N1} & y_{N1} & z_{N1} \end{pmatrix} \begin{pmatrix} 1 \\ 1 \\ 1 \end{pmatrix}$ | $\begin{pmatrix} 0 & 0 & 0 \\ 0 & 0 & 0 \\ 0 & 0 & 0 \end{pmatrix} \begin{pmatrix} 0 \\ 0 \\ 0 \end{pmatrix}$ |
| C4 (C/U only) |  |  |
| N1 (C/U only) |  |  |

When atoms were missing in a structure – which occurs frequently for the leading phosphate group – the corresponding coordinate and mask values were set to zero. The two atoms that determined the position of the phosphate backbone (first two rows in coordinate matrix) are shared between purines and pyrimidines. While both Purines and Pyrimidines have C2-atoms, this atom does not occur at the same position in the nucleobases’ ring structures and hence was encoded separately (see red frame in Fig. 1).

#### Distance and mask tensors

For an RNA sequence of length *L*, we computed for all of the *L* × *L* possible residue pairs the Euclidian distances between the encoded atoms. The resulting 8*L* × 8*L* distance matrix was re-shaped into a *L* × *L* × 64 distance Tensor *D* and symmetrized to satisfy the symmetry condition *D_ijk_* = *D_jik_*. For example, *D*_1*L*1_ = *D*_*L*11_ is the Euclidian distance between the phosphate atoms of the first and the last residue.

Some atom-atom distances in this distance tensor *D*, however, did not correspond to any meaningful values because the corresponding atoms were missing and thus the coordinate entries in the 8 × 3 matrices were zero. The respective elements in *D* were set to zero by multiplying distance tensor *D* with a mask tensor *M* that was calculated as follows. First, the mask arrays of size 8 × 1 (Supplementary Table 1) for the *L* residues were stacked to a single array of size 8*L* × 1 and the outer product with itself was calculated to obtain a matrix of size 8*L* × 8*L*. This matrix was then reshaped into a tensor of size *L* × *L* × 64 and symmetrized to obtain the mask tensor *M*.

#### Structures with long or multiple chains

When a structure had multiple chains or a chain’s length exceeded 100 nt, we carried out the following steps to obtain multiple, smaller substructures suitable for model training. First, when a structure had multiple chains, a chain was randomly selected with probability proportional to its length and used as a substructure. Then, if that chain’s length exceeded 100 residues, a random, continuous subsection of that chain was cut out and used as the substructure instead. For each residue in this substructure, the distances to the residues of the remaining, overall structure were calculated. For distances below a threshold of 3.3 Å, the corresponding residues of the substructure were flagged as “fixed” and the corresponding distance classes between “fixed” residues were presented at the input during training of the generator.

#### Data augmentation

SimRNA is a 3D RNA structure prediction software that makes use of coarse-grained residue representations and Molecular Dynamics Methods to sample the conformational space ^[7]^. This program starts with a circular initial RNA structure (that resembles a snake biting it’s own tail) and then folds the RNA to minimize an energy function while slowly cooling down the thermodynamic system. To do data augmentation, we used SimRNA the opposite way: we started with the original PDB structure and then increased the temperature to “drift away” from the original structure. Using this method, we generated 100 of such “drift structures” for each training example that were approximately 1, 3, 5 and 10 Å RMSE away from the original PDB structure.

#### Simulated SHAPE data

SHAPE (selective hydroxyl acylation analyzed by primer extension) ^[26]^ is a method for obtaining RNA secondary structure information. SHAPE exploits the fact that RNA residues that do not engage in base pairing, such as residues in dangling ends or loops, react more easily with certain reagents, making their detection possible by primer extension. In other words, higher SHAPE reactivities indicate regions of higher RNA flexibility. SHAPE data generated under comparable experimental conditions is only available for a very limited number of PDB structures ^[26]^. We hence used above mentioned drift data (see “Data augmentation”) to simulate SHAPE data. Specifically, we structurally aligned original PDB structures and their drift structures and used the average absolute position-specific deviation (that is, the RMSE between individual atoms) as simulated SHAPE data.

#### Experimental SHAPE data

We used publicly available SHAPE data^[26]^. As the simulated SHAPE data only correlates with but does not exactly match the experimental SHAPE data, we rescaled the experimental SHAPE data. For this purpose, we mapped all data percentiles between experimental and simulated SHAPE data (for example, a SHAPE value that fell into percentile 10 among the original SHAPE data was mapped to the 10^th^ percentile of the simulated SHAPE data). This mapping was learned using the following PDB IDs: 2*L*1*V*, 2*K*95, 3*PDR*, 3*DIG*, 1*P* 5*O*, 3*G*78 and 2*N* 1*Q*.

#### Similarity clustering

Structure is generally more conserved than sequence, hence structural similarity should ideally be used to split the data set into training, test and validation data sets that share as little homology as possible. However, similarity scores based on RMSE have the problem that global rearrangements dominate this score even if all structural domains are conserved perfectly. We therefore decided to use sequence similarity as a proxy for structural similarity and hence functional similarity. We aligned all sequences of all chains in the data set with a scoring function that rewarded matches with 1 and punished mismatches and opening gaps with −1. Gap extensions were treated as neutral. Then, we summed up the scores for structures that had multiple chains, and divided the scores by the overall length of the longest of each compared sequence. Using these length-specific matching scores, we used complete hierarchical clustering and chose a reasonable cutoff (0.7) to obtain sequence clusters.

#### Data Sampling

First, we sampled uniformly over all sequence clusters. Longer sequences are more informative than shorter sequences, hence we sampled proportional to sequence length.

#### Homologs

To obtain sequences homologous to those in our PDB structure data set, we followed the following workflow. First, we downloaded homologous sequences from RNAcentral ^[29]^ using a Python script published on the RNAcentral Sequence search API website ^3^. Then, we only kept homologous sequences that fulfilled the following conditions (i) their E-value had to be less than 0.01, (ii) they must not have insertions, and (iii) they had to have at least one mutation other than a deletion. Gaps in homologous sequences were filled up with the letter ‘N’.

### Deep Learning Architectures and Hyperparameters

#### VQ-VAE Network Architecture

We encoded RNA tertiary structures by distance tensors (*L* × *L* × 64) – as explained above – that act as both input and target of our VQ-VAE. We zero-padded all entries of distance tensors that were outside the maximum sequence length *L* = 100. Following the original work of the VQ-VAE^[20]^, the encoder architecture was a residual network ^[30]^ that consisted of one convolutional layer followed by 4 residual blocks. Each residual block was of the form: Block = 2x[Batchnorm, ReLU Activation, Conv]. The two convolutional layers (“Conv”) within each block could have different convolutional kernels, whose sizes we report in the following format hereafter: [hight × width, out channels].

**Supplementary Table 2.**
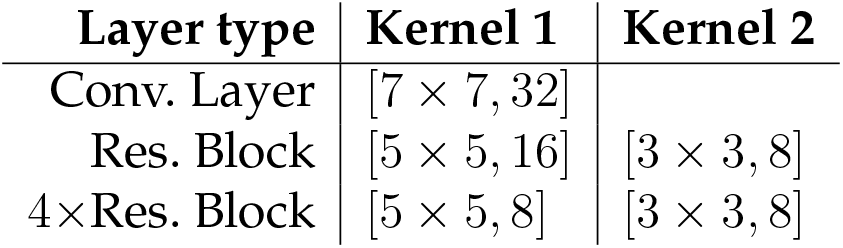
VQ-VAE encoder architecture.

In the vector quantisation step, the encoder’s residual network output of shape (*L* × *L* × 8) was mapped to VQ-VAE encodings (distance classes) of shape (*L* × *L* × 3) using 3 codebook vectors of embedding dimension 8. We enforced symmetry of the distance class tensors (along the first 2 dimensions) by adding the transpose of the embedding and dividing the result by 2. The decoder architecture took the VQ-VAE encodings (distance class tensor) as input and consisted of four sets of residual blocks followed by a final ReLU layer.

**Supplementary Table 3.** VQ-VAE decoder architecture.

| Layer type | Kernel 1 | Kernel 2 |
| --- | --- | --- |
| Conv. Layer | $[5 \times 5, 8]$ | |
| 4×Res. Block | $[5 \times 5, 8]$ | $[5 \times 5, 8]$ |
| Conv. Layer | $[5 \times 5, 16]$ | |
| 4×Res. Block | $[5 \times 5, 16]$ | $[5 \times 5, 16]$ |
| Conv. Layer | $[5 \times 5, 32]$ | |
| 4×Res. Block | $[5 \times 5, 32]$ | $[5 \times 5, 32]$ |
| Conv. Layer | $[5 \times 5, 64]$ | |
| 4×Res. Block | $[3 \times 3, 64]$ | $[3 \times 3, 64]$ |
| ReLU Activation |  |  |

Between each set of residual blocks, we up-scaled the number of feature-maps by inserting an additional convolutional layer that doubled the number of feature maps. The output of the decoder had shape (*L* × *L* × 64) and corresponded to the distance tensor reconstruction. For training, we used the same learning objective as in the original VQ-VAE^[20]^ using an exponential moving average for the codebook vector updates with a decay rate of 0.99. For optimization, we used standard Adam Optimizer^[31]^ with a learning rate of 1 × 10^−5^ at a batch size of 100.

#### Generator Network: Data Preprocessing

We stacked the following 4 tensors to obtain input tensors for the generator network: (i) an encoded RNA sequence tensor, (ii) the corresponding, partially masked distance class tensor, (iii) coordinate frame tensors to provide the network with positional information, and (iv) an optional attention map of homologous sequence alignment and SHAPE data. RNA sequences of length *L* were encoded as unique bit patterns of shape (*L* × *L* × 8), (see Supplementary Fig. 1). First, an RNA sequence was one-hot encoded by a tensor of shape (*L* × 4). Unknown nucleotides that were denoted by an “N” in the sequence were encoded by setting all values in the one-hot encoding to 0.25. This one-hot encoded sequence was then copied *L* − 1 times to obtain a tensor of shape (*L* × *L* × 4). Then, this tensor and its transpose were stacked along the last dimension to obtain a tensor of shape (*L* × *L* × 8). This tensor corresponded to a unique bit pattern for each possible pairing and also contained directional information. For sequences with *L* < 100, the sequence tensors were uniformly padded with −1.

Partially masked distance class tensors were obtained from the distance classes using a pre-trained VQ-VAE, followed by a “partial masking” process in which some pixels were set to zero as described in the following. First, we drew the number of pixels to be masked at a given training step from a truncated normal distribution with mean *L*^2^/2, standard deviation *L*^2^/4 and which was bounded at 2 standard deviations around the mean (this ensured that at least 0 and at most *L*^2^ pixels were masked, while rarely masking either very few or very many pixels). Then, we randomly drew the corresponding number of pixel positions and encoded masked pixels with (0, 0, 0). However, we found that encoding masked pixels this way made it difficult for the network to learn the difference between masked distance classes (0, 0, 0) and the regular, non-masked distance classes (1, 0, 0), (0, 1, 0) or (0, 0, 1). To overcome this limitation, we encoded all regular, non-masked distance classes by setting all zeros to −1, so that, for example, (1, 0, 0) was encoded as (1, −1, −1). Coordinate frames consisted of a “diagonal frame” that had ones along the diagonal and was zero elsewhere, and a “padding frame” which contained a box of side length *L* that had ones along its border and was zero elsewhere. The coordinate frames were padded with −1 when sequences were shorter than 100 nt. The production of the attention map of homologous sequences and SHAPE data is described in the following section in detail. Overall, the generator input was a tensor of shape (*L* × *L* × 14), and contained the encoded sequence tensor of shape (*L* × *L* × 8), the partially masked distance tensor of shape (*L* × *L* × 3), both coordinate frames of shape (*L* × *L* × 2) and the homologous sequences alignment and SHAPE data attention map tensor of shape (*L* × *L* × 1).

#### Generator Network: Attention map of homologous sequences and SHAPE

For each training example, an array of 50 randomly chosen, aligned and one-hot encoded homologous sequences was produced. While the 4 standard nucleotides were encoded using one-hot coding (e.g. “A” was encoded as (1, 0, 0, 0)), gaps and unknown nucleotides were encoded by setting all possible one-hot coding values to 0.25 such that the one-hot coding vector became (0.25, 0.25, 0.25, 0.25). When there were fewer than 50 homologous sequences, the original sequence was used to “fill up” the array. The resulting array of one-hot encoded homolgeous sequences had shape (*L* × 50 × 4). Using a standard dense layer, this array was mapped onto a tensor of shape (*L* × 50). Then, simulated SHAPE data was included by stacking a vector containing *L* SHAPE reactivity values on that array, resulting in a tensor of shape (*L* × 51). Using two separate dense layers, two tensors of shape (*L* × 64) were produced and self-attention was applied to these tensors by using them as query and key tensors as described in the original Transformer paper^[18]^, resulting in an attention map of shape (*L* × *L* × 1). Self-Attention added the benefit, that for every pixel, the corresponding nucleotide could receive both homologous sequence information as well as SHAPE reactivity information from all other nucleotides in that sequence. We computed results on a sample attention map for the Adenine Riboswitch (PDB: 1y26) in Supplementary Fig. 2.

#### Generator Network: Architecture

Using the (*L* × *L* × 14) input tensor described above, the generator network performed regression on the full distance class map under the masked learning objective, which is specified further below. The input tensor was first passed through a convolutional layer to be further processed by a deep residual network architecture with eight blocks, each one having the following structure: ResBlock = 2x[Batchnorm, Elu Activation, Conv] + 1x[Batchnorm, Elu Activation, Conv] + 1x[Batchnorm, Elu Activation, DilatedConv], with standard skip connections after the first two and between the last two subblocks. We employed standard convolutions with kernel sizes as described in Supplementary Table 4 below. We also used dilated convolutions with a dilation rate of 2 in the residual blocks. Here, the last convolution layer compressed the residual network output into an appropriate shape of (*L* × *L* × 3) so that, after a final softmax activation, the network’s output corresponded to predicted distance classes.

**Supplementary Table 4.**
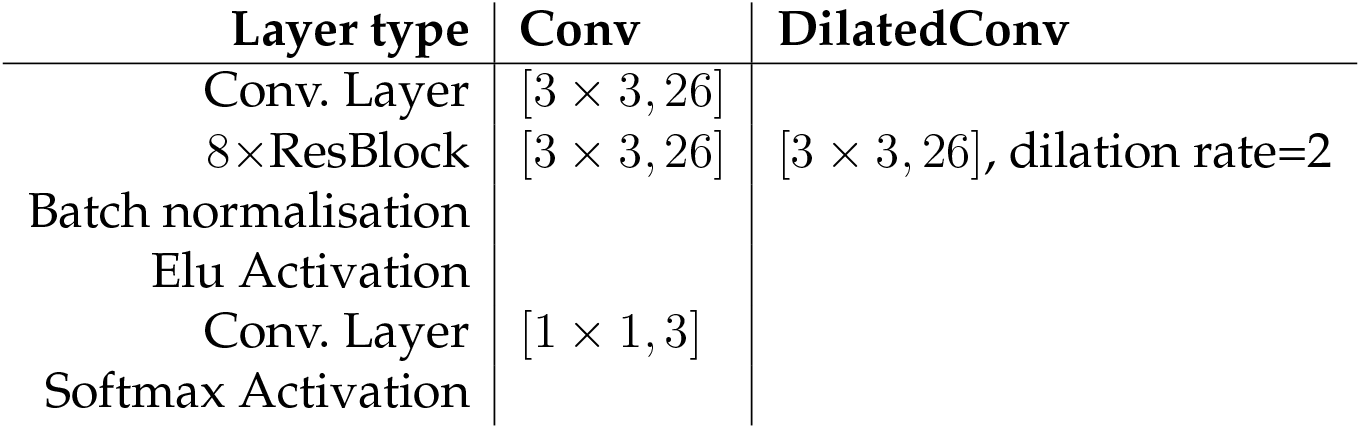
Generator network: Resiudal architecture.

The network was trained under the masked learning objective, implemented using the cross entropy between target distance classes and generator predictions coming from input tensors with partially masked distance tensors as a loss function. We further employed weight decay (*L*_2_ regularization) following standard recommendations for training large residual architectures^[30]^. We used the following hyperparameter setup using a lazy Adam Otimizer.

**Supplementary Table 5.** Generator network: Optimization hyperparameters.

| LazyAdamOptimizer | $\beta_1 = 0.9, \beta_2 = 0.997, \epsilon = 1e - 8$ |
| --- | --- |
| Batch size | 500 |
| Weight decay | 0.01 |
| Learning rate $\alpha$ | 0.001 |
| Learning rate warmup steps | 100000 |

For stabilized training, we chose a learning rate with linear warmup scaling and cosine decay. After two million iteration steps we stopped the network training.

#### Score Model

The Score Model was implemented to distinguish which one of two distance class maps was better. Its architecture was a residual network as described in the following table.

**Supplementary Table 6.** Score Model network architecture.

| Layer type | Conv | DilatedConv |
| --- | --- | --- |
| Conv. Layer | $[3 \times 3, 5]$ | |
| 4×ResBlock | $[3 \times 3, 5]$ | $[3 \times 3, 5]$, dilation rate=2 |
| Batch normalisation |  |  |
| Relu Activation |  |  |
| Conv. Layer | $[1 \times 1, 100]$ | |
| Relu Activation |  |  |
| Conv. Layer | $[1 \times 1, 1]$ | |

The Score Model was trained to discriminate between “correct” and “incorrect” distance class maps as described in the following. The set of correct examples consisted of all 8048 examples in the training data set. To create the set of incorrect examples, we used SimRNA-based data augmentation to compute 100 drift structures of up to 10 Å RMSE away from each original structure in the training data set. Using the VQ-VAE setup, we then computed distance class maps for all original as well as drift structures. We further increased the dataset of incorrect examples by factor 10 by randomly flipping class pixels from the targets distance classes. We also used the Generator networks’ argmax predictions, showing only 0%, 5%, 10%, 15% and 20% of the target distance classes at the input. Then, at each training step, both a correct and a corresponding incorrect distance class map of shape (*L* × *L* × 3) were fed through the network to obtain two “logit maps” of shape (*L* × *L* × 1). The values of the correct logit map were then subtracted from corresponding values of the incorrect logit map, followed by taking the sum over all differences. We optimized this objective with *L*_2_ regularization under the following hyperparameter setup and used a standard learning rate decay exponential to the iteration steps.

**Supplementary Table 7.**
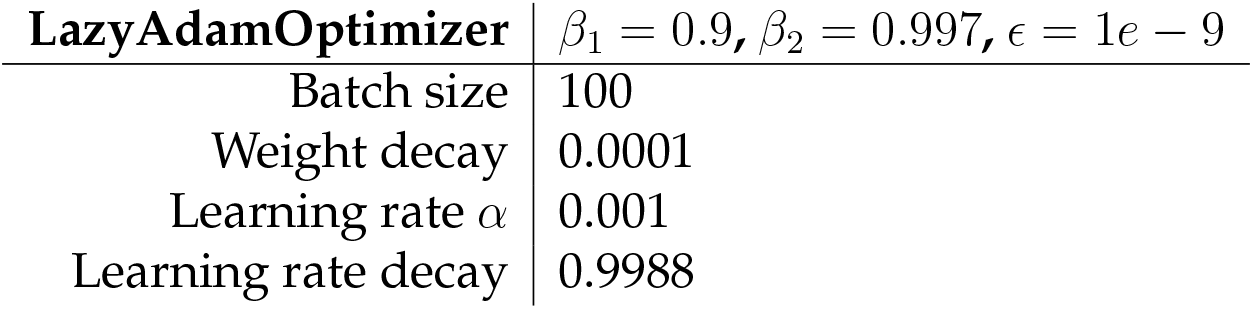
Score Model network: Optimization hyperparameters.

| **LazyAdamOptimizer** | $\beta_1 = 0.9, \beta_2 = 0.997, \epsilon = 1e - 9$ |
| --- | --- |
| Batch size | 100 |
| Weight decay | 0.0001 |
| Learning rate $\alpha$ | 0.001 |
| Learning rate decay | 0.9988 |

#### MCTS: Sampling structural ensembles

The MCTS algorithm sampled pixelwise one of the three distance classes using the Generator network iteratively. Naturally, the diagonal had high class probabilitiy values, so that we started initialising the search tree by setting the diagonal to the nearest distance class. Except for the diagonal, we started by masking all (*L* × *L* × 3) target distance classes with zeros and iteratively filling in class indicators, e.g. (1, −1, −1) if the MCTS chose the first distance class. With that basic step logic, MCTS could iteartively fill up pixels and update visit counts and values at each node. The value objective in the MCTS was designed such that the network aimed to get a sharp view after some pixels were set, so that not every single of the *L*^2^/2 pixels needed to be sampled. This reduced the depth of the search tree by a large margin. Typically, the Generator network produced sufficiently sharp predictions when the MCTS was able to predict 30% of the target distance classes correctly. The remaining pixels were then filled up using an argmax prediction of the Generator network given that leaf. Exemplarly, we computed the leafs of the search tree for two alternative structures shown in Supplementary Fig. 3for the ZMP-Riboswitch (PDB: 4xw7).

---

**Supplementary Figure 1.**
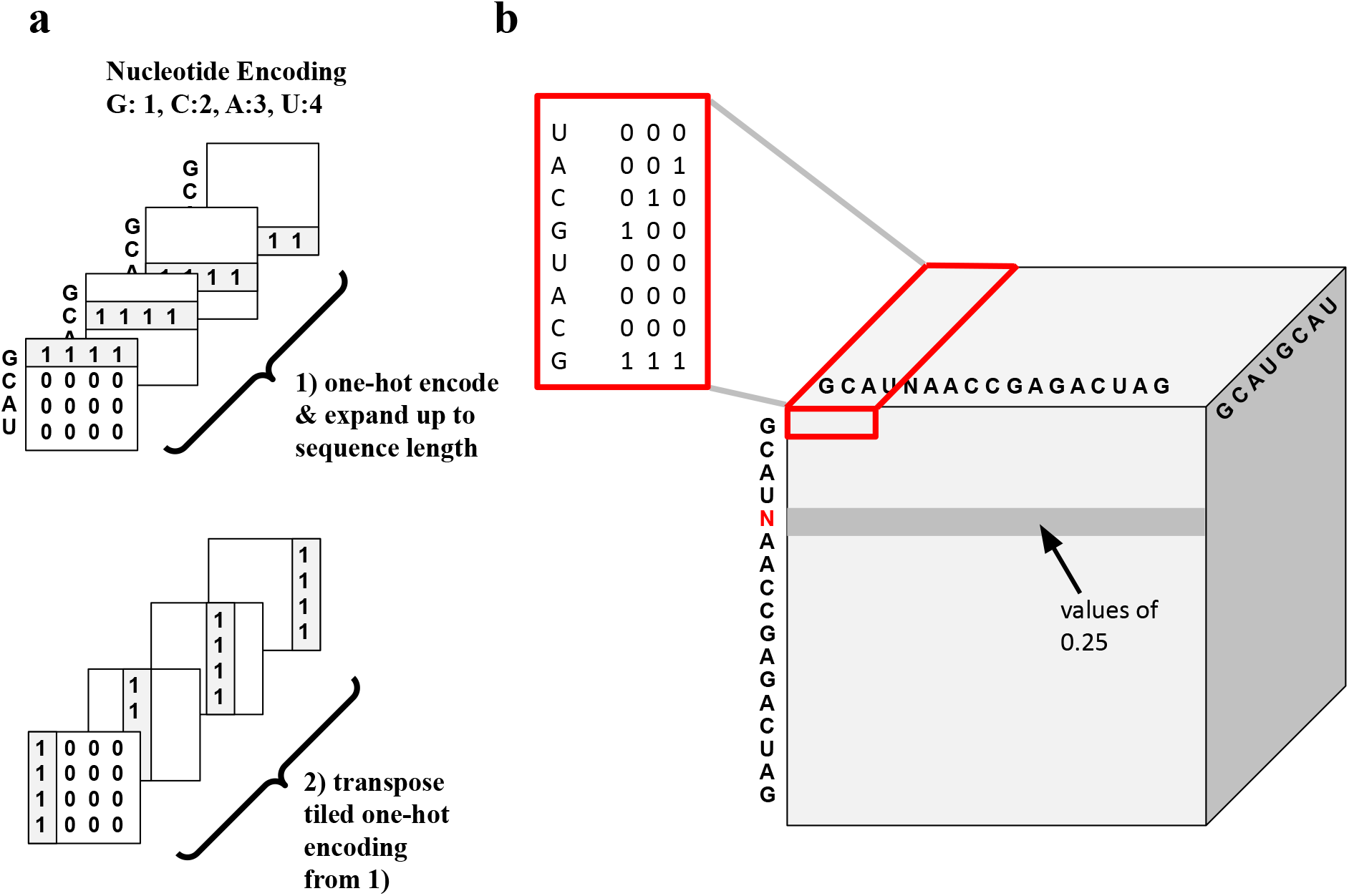
Encoding of Sequence Information. RNA sequences of length *L* were encoded as unique bit patterns of shape (*L* × *L* × 8): **a** First, every nucleotide in an RNA sequence was one-hot encoded, e.g. *G* : (1, 0, 0, 0), *C* : (0, 1, 0, 0), *A* : (0, 0, 1, 0), *U* : (0, 0, 0, 1), *N* : (0.25, 0.25, 0.25, 0.25). For the full sequence, these one-hot encodings led to a tensor of shape (*L* × 4). Unknown nucleotides that were denoted by an “N” in the sequence were encoded by setting all values in the one-hot encoding to 0.25. This one-hot encoded sequence was then copied *L* times (a1) to obtain a tensor of shape (*L* × *L* × 4). Then, this tensor and its transpose (a2) were stacked along the last dimension to obtain a tensor of shape (*L* × *L* × 8). **b** A sample sequence Tensor that corresponded to a unique bit pattern for each possible pairing and also contained directional information. For sequences with *L* < 100, the sequence tensor was uniformly padded with −1 Red insert: example bit pattern for the Tensor at the first three pixels with depth 8.

**Supplementary Figure 2.**
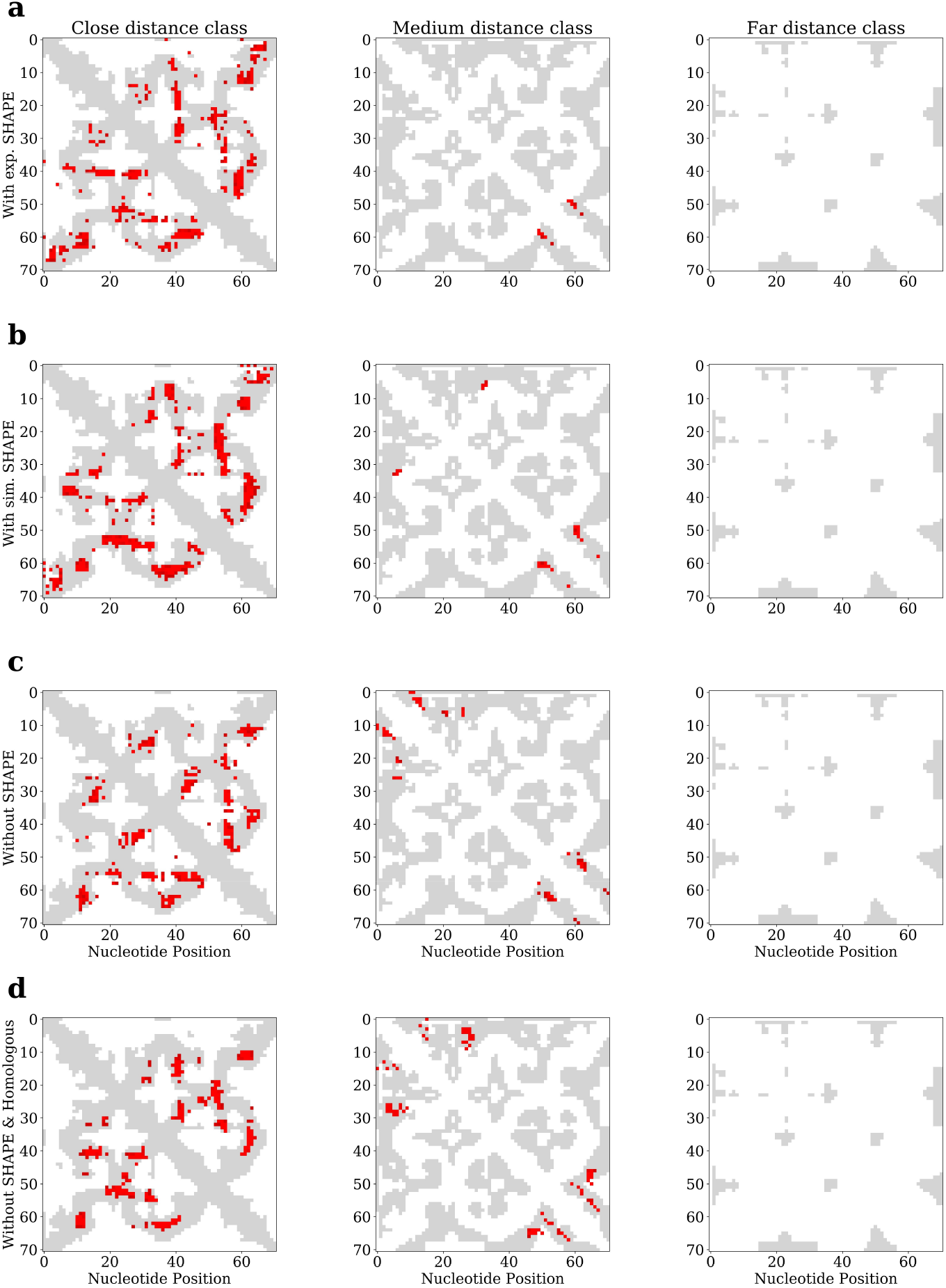
Attention: Improved performance from adding Structural Probing data (SHAPE) and homologous sequences. Attention Maps were a good indicator for the location of structural contacts. High attention values (red pixels) were almost exclusively found at the “near” distance class, resulting in higher probability scores for the generator’s initial prediction for that class. Incorporation of experimental **a** or simulated **b** SHAPE data both resulted in lower false positive rates of attention placement compared to when no SHAPE data **c** or no SHAPE data and no homolgous sequence information **d** was used, with false positive rates being, 4.3%, 5.7%, 14.8% and 21.4%, respectively (the false positive rate was calculated as the number of incorrectly placed red, high attention points divided by the total number of red, high attention points, where high attention points were defined by attention scores above 0.01.)

**Supplementary Figure 3.**
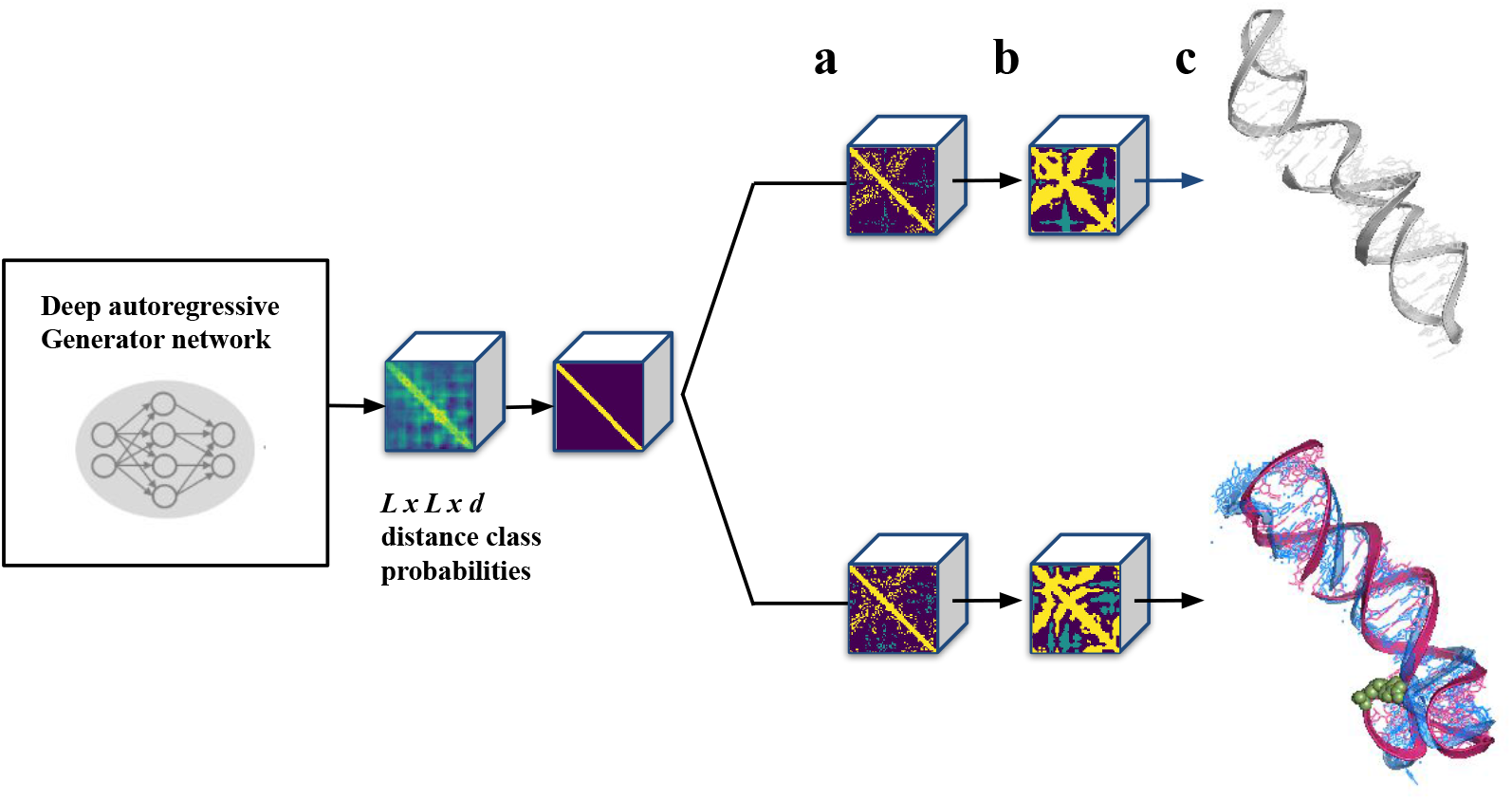
MCTS: Search tree with two distinct leafs. Starting with all target distance classes masked, the Generator places initial probabilities for every pixel in the distance class softmax prediction. From there, pixels were sampled iteartively using the MCTS search objective which aimed for entropy reduction. We derived two terminal leaf nodes **a** for which the Generator network saw enough distance pixels to fill up the remainers using its argmax prediction **b**. From those filled up leafs, the VQ-VAE could be applied to decode into real distance space **c**. After further energy refinement, we showed for the ZMP-Riboswitch (PDB: 4XW7), that those two leafs indeed corresponded to two different structures. The riboswitch has a movable hinge part, which gets stabilized by a small molecule. Hence, our best leaf prediction in red is closer to the blue target solution, bottom **c**. We also sampled an alternative structure. In this particular example, the strechted grey RNA structure **c** was ranked with a lower score by the Score Model

## Footnotes

1 https://files.rcsb.org/pub/pdb/derived_data/pdb_entry_type.txt

2 http://files.rcsb.org/download/

3 https://rnacentral.org/sequence-search/api

